# Scalable variant-to-function screens using prime editing perturb-seq

**DOI:** 10.64898/2026.10.28.762453

**Authors:** Joachim De Jonghe, Magdalena Armas Reyes, Catriona Jamieson, Jens Bager Christensen, James W. Opzoomer, Jake P. Taylor-King

**Affiliations:** Relation, London, United Kingdom; Gurdon Institute, Cambridge University, Cambridge, United Kingdom; Department of Physiology, Development and Neuroscience, Cambridge University, Cambridge, United Kingdom

## Abstract

Linking genetic variants to their functional consequences at scale remains a major challenge in human genetics. Perturb-seq is limited in the variants it can install, while prime editing has largely been restricted to simpler mechanistic readouts. Here, we introduce prime editing perturb-seq (PEP-seq), which links programmed mutations to transcriptomic states. We designed 2,164 prime editing guides targeting common expression quantitative trait loci (eQTLs), premature termination codons, and position-matched synonymous controls, generating 749,110 time-resolved single-cell transcriptomes. PEP-seq precisely detected nonsense-mediated decay, while probe-based junction measurements suggested depletion of mature RNAs. For eQTLs, a locus-size-corrected specificity metric showed that the intended target was frequently the most affected transcript within a 1 Mb window, with effects also associated with open chromatin near the edit site. These results establish PEP-seq as a scalable platform for direct, endogenous variant-to-function mapping.

## Introduction

Genome-wide association studies and molecular QTL mapping have identified hundreds of thousands of candidate regulatory variants, but distinguishing causal variants from their correlated neighbors, and establishing which gene(s) and cellular phenotypes a variant perturbs requires direct functional interrogation^1,2^. Massively parallel reporter assays (MPRAs) enable thousands of variants to be tested simultaneously for effects, but typically assay short sequences outside their native genomic context, where splicing, chromatin state and long-range regulatory interactions are not preserved^3,4^. Consequently, reporter activity does not necessarily establish an accurate representation of a variant’s effect on cellular phenotypes. Assaying programmed variants genome-wide remains challenging: CRISPR/Cas9 can introduce engineered variants, but the requirement for guide RNA and homology-directed repair necessitates assaying variants in separate compartments^5^, lowering scale. Base editing has enabled much more scalable installation of variants directly in the endogenous genome without double-strand breaks or donor templates. However, base editors remain constrained by the chemistry of the available nucleotide conversions, PAM availability and editing-window position, and can introduce bystander edits when multiple editable bases occur within the activity window^6^. Thus, only a subset of naturally occurring variants can be modelled cleanly using base editing. On the other hand, prime editing enables precise endogenous variant installation, necessitating only PAM availability without the aforementioned pitfalls in scale and range, making it well suited to study mutation effects^7^.

However, most pooled prime-editing screens to date rely on fitness or sensor readouts^8–11^, missing more mechanistic effects on *cis-*regulation, pathway activity, cell state, and neomorphic transcriptional programs in *trans*. Coupling prime editing with single-cell RNA-seq would therefore enable a richer view of allelic function in native genomic context. Although feasibility has been demonstrated^12^, existing studies are limited in the scale and scope required to meaningfully mine large sets of variants across the genome.

Recent advances in single-cell technologies have substantially increased in throughput via combinatorial indexing of single-cell material^13,14^. Most widely-used is the 10x genomics Flex Apex solution, which captures up to 1 million cells per GEM-X well via mRNA pre-indexing. Its integration with genome-wide perturb-seq screens was further demonstrated with VIPerturb-seq^15^, but a framework for harnessing the scalability offered by these methods to screen for variant effects is currently lacking.

Here, we developed a scalable methodology to integrate pooled prime editing screens with single-cell readouts: PEP-seq. We designed a library of 2,164 prime-editing guide RNAs (pegRNAs) spanning three variant classes to test our platform: (i) candidate *cis-*eQTLs to test variant directionality and target-gene specificity; (ii) premature termination codons (PTCs) at essential and non-essential genes with position-matched synonymous controls, to measure nonsense-mediated decay (NMD) with a built-in specificity control; and (iii) a smaller, exploratory set of candidate splicing QTLs (sQTLs), to explore splicing-dependent *trans-*regulatory effects. Guides were transduced into a clonal HEK293T line constitutively expressing prime editor 7^16^ (PE7) and transcriptomes were assayed with a modified VIPerturb-seq framework to provide sufficient cellular coverage for each variant. Below we characterize the technical performance of this platform, accurately quantifying gene knockdowns via nonsense-mediated decay and recovering cis-regulatory effects of candidate eQTLs, showing a direct link between local chromatin accessibility and the probability of detecting a significant effect.

## Results

### Design and quality control of PEP-seq (Figure 1)

A modified version of VIPerturb-seq was used to enable pooled, single-cell readout of prime-editing guide libraries at scale (Fig. 1a). Prime-editing guide RNAs were designed using PRIDICT2^17^ and cloned, together with polyT termination, stuffer and right-hand side (RHS)/left-hand side (LHS) barcode sequences into a CROP-seq^18^ lentiviral backbone (Supplementary File 1). The library was transduced into triplicate plates of HEK293T cells constitutively expressing PE7^16^ at a multiplicity of infection (MOI) of 0.1, so that the great majority of transduced cells receive a single integrated guide. Cells were selected for guide integration by puromycin selection and harvested at days 4, 10 and 17 post-transduction, fixed, and hybridized with both the standard human transcriptome probe set and custom VIPerturb-seq guide-identity probes before combinatorial fluidic indexing^19^ on the GEM-X chip for paired gene-expression and guide-capture library preparation.

**Figure 1.**
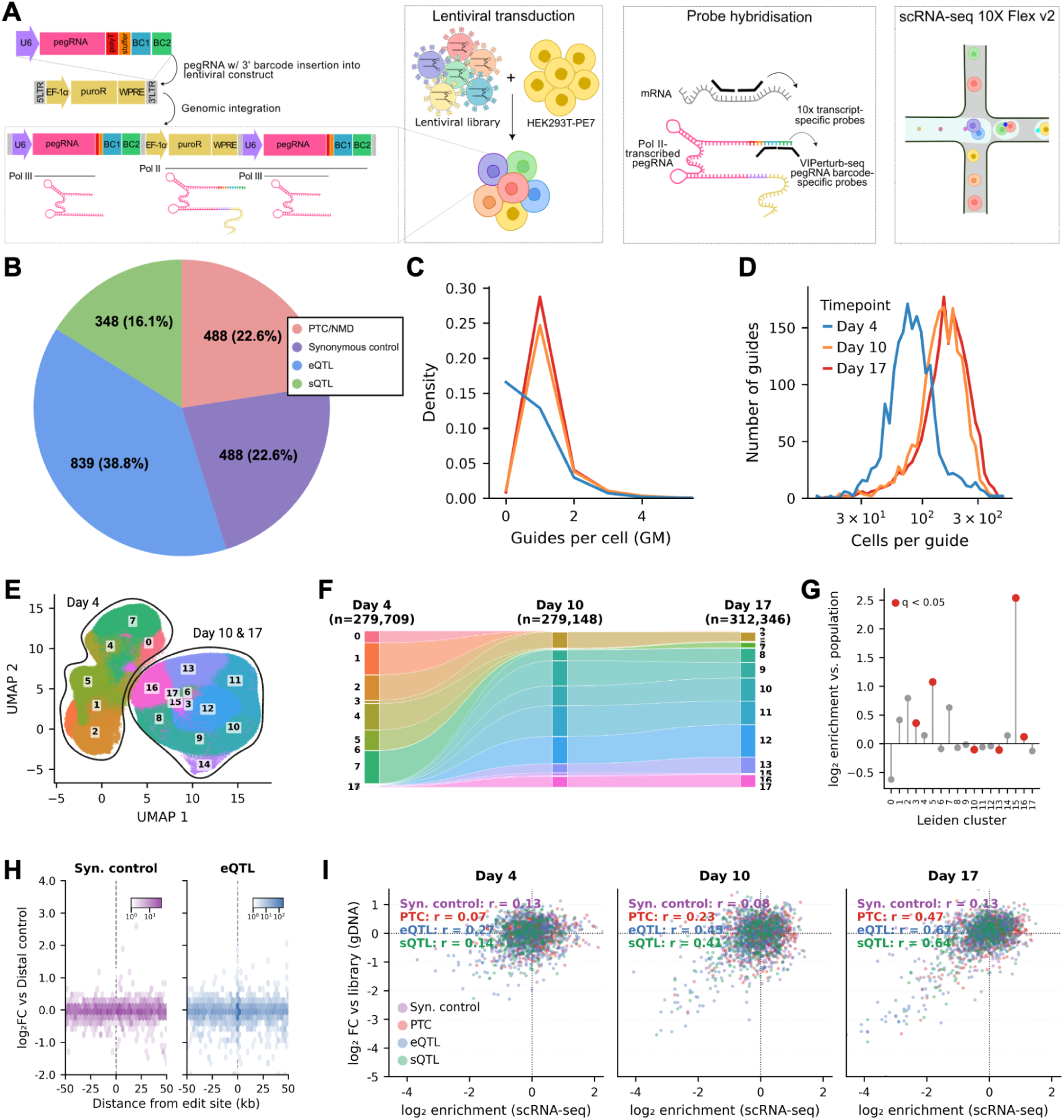
Design and quality control of pooled prime editing screen with single-cell readouts using probes **(a)** Schematic of the pooled prime-editing screen: pegRNA guide libraries, together with polyT, stuffer and VIPerturb-seq RHS/LHS barcode sequences, are cloned into a CROP-seq lentiviral backbone and delivered at MOI 0.1 to HEK293T cells stably expressing prime editor 7 (PE7). Cells are harvested at days 4, 10 and 17 post-transduction, fixed, and hybridized with human transcriptome and custom guide-identity probes before single-cell processing. **(b)** Composition of the 2,164-guide pegRNA library by variant class: 839 candidate cis-eQTLs (38.8%), 488 premature-termination-codon (PTC) edits (22.6%) and their 488 position-matched synonymous controls (22.6%), and 348 candidate splicing-QTL (sQTL) variants (16.1%) arising from a post-hoc re-audit of the eQTL candidate pool (Methods); one residual “Other” design completes the panel. **(c)** Distribution of guides detected per cell (guide multiplicity, GM) at each of the three post-transduction timepoints as calculated on 749,110 post-QC cells (with low quality guides excluded). **(d)** Guide representation across the pegRNA library at each timepoint: number of distinct guides (of 2,164 total) reaching a given number of QC-passing cells, same population as (c). **(e)** UMAP embedding of 871,302 QC-passing cells (low quality guides included) coloured by Leiden cluster (resolution = 1.0; Methods; 18 clusters, 559–116,157 cells each). **(f)** Sankey diagram of the same 18 Leiden clusters’ relative composition across the three timepoints (Day 4: n = 279,709; Day 10: n = 279,148; Day 17: n = 312,346). **(g)** Enrichment of the 18 Leiden clusters (e) among guides with a strong bulk fitness effect (|gDNA log2 fold-change vs. plasmid library| > 1 at day 17; Methods), tested per cluster against the day-group population baseline (one-sample binomial test, BH-corrected across clusters; day 10+17 combined). **(h)** Hexbin density of log2 fold-change (guide vs. distal synonymous-control background, days 10 and 17 pooled, target genes excluded) against signed distance from the edit site, for all guides across all four variant classes with sufficient cell numbers (n = 8,794 guide–gene pairs, restricted to ±50 kb of the edit site for display). **(i)** Cross-validation of guide representation between single-cell CRISPR-capture (scRNA-seq) and bulk genomic-DNA (gDNA) amplicon sequencing of the same lentiviral pegRNA pool, for 1,827 pegRNA designs recovered in both assays: per-class Pearson correlation between scRNA-seq cell-count enrichment and gDNA fold-change (vs. the day-0 plasmid library), by timepoint.

To assess if the workflow would enable linking variants to their single-cell transcriptome, we designed a library of 2,164 pegRNAs targeting 1,533 distinct genomic positions, with up to three independently designed pegRNAs per site, and divided into four variant classes (Fig. 1b): 839 candidate cis-eQTLs (38.8%), 488 premature-termination-codon (PTC) edits (22.6%) with 488 position-matched synonymous controls (22.6%), and 348 candidate splicing-QTL (sQTL) variants (16.1%). eQTLs and sQTLs were prioritised from the EMBL eQTL catalogue^2^, based on effect size and directionality, frequency across datasets and PRIDICT2^17^ editing scores.

Knockdown targets via NMD in HEK293T cells were prioritized from the X-atlas/Orion data^20^ based on large effect size and PRIDICT2 editing scores, targeting 5’ tyrosines to introduce either a premature termination codon or a matched synonymous variant (see Methods), as previously detailed by Cirincione *et al^10^*.

After single-cell quality control (retaining cells with 5,000-75,000 total UMI counts and ≤5% mitochondrial reads), 871,203 of 1,200,369 sequenced cells (72.6%) passed QC. To further validate library integrity, genomic DNA pellets for each timepoint were collected and sequenced along the plasmid library; construct-integrity filtering (see Methods) removed a further 122,093 cells (10.3%,15.9%, and 15.6% at days 4, 10, and 17, respectively), leaving 749,110 cells for final analysis (Supplementary Fig. 1a,b). At day 4, 49.5% of the cells carried no detectable guide, indicating ongoing puromycin selection. However, selection was mostly achieved by day 10 and day 17, with the fraction of cells without a called guide falling to 3.8% and 2.5%, respectively (Fig. 1c). The large majority of the cells in the dataset only had one guide assigned (66.3% of all cells), indicative of successful transduction and selection at a MOI of 0.1. The cellular coverage per guide follows the same trajectory, rising from a median of 90 cells/guide at day 4 to 162 at day 10 and 177 at day 17 across the entire guide library (Fig. 1d).

A UMAP embedding shows cells at day 4 occupying a distinct, transcriptionally heterogeneous region of the embedding, consistent with an early selection stage, while day-10 and day-17 cells converge on states that are essentially unchanged between the two later timepoints; this was demonstrated by Leiden clustering followed by assessing which time points were represented within each cluster (Fig. 1e,f). By plotting other key QC statistics over the UMAP coordinates, we investigate if confounding factors have skewed our analysis. Replicates appear reproducible and differences in pseudobulk gene expression profiles suggest the cell transcriptome stabilizes at later time points (Supplementary Fig. 1c). Investigating the number of guides, total UMI, mitochondrial counts, and cell cycle status (using a rank-based method) suggests no obvious confounding biasing the experiment (Supplementary Fig. 1d); in addition no guide is enriched for any particular cell cycle state (no guide reaching FDR significance; Supplementary Fig. 1e) as has been noted in the analysis of other perturb-seq experiments^21,22^.

We next asked whether guides perturb a cell’s overall transcriptional state. Restricting to cells with multiplicity of one, for every guide with ≥25 cells identified we tested whether its cells’ cluster distribution departs from the population baseline via Jensen-Shannon divergence (JSD) against a matched-n permutation null (Methods). Using cells harvested at either day 10 or day 17, we see that a group of guides induce cells to concentrate specifically in cluster 15 (0.36% of the atlas), at up to ∼5.8-fold the expected rate (Fig. 1g). The markers for cluster 15 include heat-shock genes (*HSPA1B*, *HSPA6*, *HSP90AA1*, *DNAJB1*, *HSPH1*), *JUN*, and mitochondrial OXPHOS transcripts (*MT-ATP6*, *MT-CO1/2*, *MT*-*ND4*) suggesting a cell-stress state.

Cas9 occupancy can interfere with transcription directly and, when targeted to cis-regulatory elements, disrupt transcription-factor binding or enhancer activity, potentially affecting genes regulated at a distance^23–25^. Because PE7 is constitutively expressed in our cell line, we asked whether persistent engagement at the target locus could modulate transcription. By examining both synonymous edits and eQTL edits (when compared to other synonymous edits targeting DNA regions more than 1Mb away), we saw no systematic reduction in expression of either the target gene or other genes within a 1Mb window; a zoom in of the nearby 50 kb window is shown (Fig. 1h). Together, these analyses indicate that prime-editor targeting does not produce detectable repression of either the directly targeted gene or neighbouring genes, positioning PEP-seq as a well-suited platform for variant-to-function studies without systematic transcriptional bias from Cas9-based targeting machinery.

To assess whether targeted genes show evidence of fitness effects consistent with essentiality, we compared each guide’s representation in sample genomic DNA (relative to the plasmid library) with its representation in single-cells with a unique guide (relative to the mean synonymous-control population) across 1,827 QC-passing pegRNA designs recovered in both assays (Fig. 1i). The two orthogonal assays are weakly correlated at day 4 for every variant class, but concordance strengthens substantially by day 10 and further by day 17 specifically for the three variant-bearing classes (eQTL, PTC/NMD, sQTL), while synonymous controls remain uncorrelated across all three timepoints (r ≈ 0.13).

Finally, these fitness effects help us understand why some guides induce changes in the distribution of cells across Leiden clusters. By splitting guides by bulk gDNA fitness effect (|log2FC| > 1 at day 17 vs. plasmid library), we see that high-fitness-effect guides exceed the null more often than low-fitness-effect guides (15/143, 10.5% vs. 132/1,987, 6.6%; p = 0.0055, 0.00075; Supplementary Fig. 1f). Guides driving a growth/fitness phenotype thus disproportionately push cells into this stress cluster, rather than perturbation type predicting departure from baseline.

Together, these results show that PEP-seq was well calibrated, yielding stable and reproducible measurements with little evidence of systematic bias from guide assignment, cellular state or prime-editor targeting.

### Detection of gene expression changes induced by nonsense-mediated mRNA decay (Figure 2)

To test whether PEP-seq could resolve subtle transcript-level effects, we used programmed premature termination codons (PTCs) as a benchmark, where precise editing is expected to produce a defined reduction in transcript abundance through nonsense-mediated decay.

**Figure 2.**
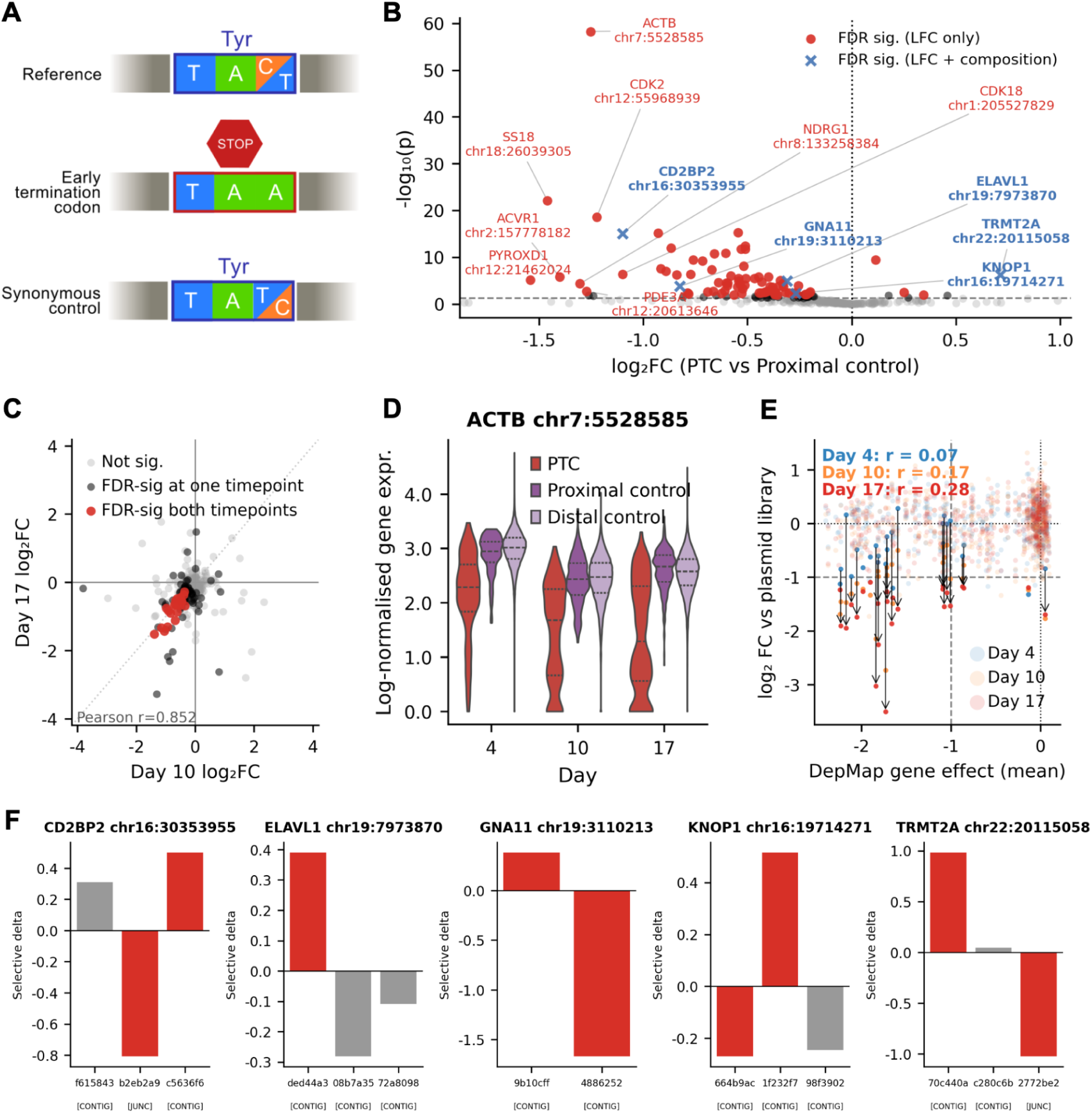
Gene expression changes induced by nonsense-mediated mRNA decay **(a)** Schematic of the PTC/synonymous-control pegRNA design: for each of 488 target genes, up to 3 candidate PTC sites (restricted to positions predicted to trigger nonsense-mediated decay; Methods) are each paired with a synonymous control edited at the same nucleotide position wherever possible. **(b)** Volcano plot of the day 10+17-combined differential expression test for each of 356 tested PTC guide positions (225 target genes) against its position-matched proximal synonymous control. Blue markers denote FDR-significant positions with corroborating probe-composition evidence (panel f). **(c)** Day 10 vs. day 17 concordance of PTC-induced log2 fold-changes, for the 339 positions tested independently at both timepoints. **(d)** Representative example of PTC-induced knockdown at *ACTB* (chr7:5,528,585), showing log-normalised expression per cell across days 4, 10 and 17, split by PTC guide, position-matched proximal control, and locus-independent distal control. **(e)** PTC/synonymous-control guide representation in bulk genomic DNA (log2 fold-change vs. the day-0 plasmid library) plotted against each guide’s target gene’s DepMap CRISPR essentiality score (mean gene effect), for 475 guide designs present at each of days 4, 10 and 17 (colour). **(f)** Probe-level selective-usage test for the 67 (of 82) FDR-significant NMD hits with sufficient probe-level coverage, testing whether transcript loss is spread evenly across a gene’s Flex probes.

Position-matched synonymous edits were included as local controls to distinguish the effect of the programmed sequence change from locus-specific background.

Our library included guides for 488 PTC edits covering 266 genes (up to 3 pegRNAs per gene targeting different tyrosine residues), each paired with a “proximal” synonymous control edited at the same nucleotide position (Fig. 2a; Methods). Post quality control (requiring ≥25 cells per guide), 356 PTC guide positions were tested against the corresponding proximal synonymous control population using cells from both day 10 and day 17 to identify 82 positions (69 genes, 23.0%) with a statistically significant change (FDR q < 0.05) in target-gene expression, of which 78 (95%) are downregulated (Fig. 2b) as expected due to nonsense-mediated decay (NMD).

The joint analysis of day 10 and day 17 was motivated by the highly reproducible effects between timepoints: by examining individual differential gene expression changes at both day 10 and day 17 separately, 21 (of 339 positions) reach significance at both timepoints, and their day 10 and day 17 log2 fold-changes are strongly correlated (Pearson r = 0.85, p = 9.4 × 10⁻⁷), and all 21 are downregulated on both days with no sign reversals (Fig. 2c). A representative example considering a guide targeting *ACTB* illustrates the time course of this knockdown, with increasing separation from the control emerging over time; this could be explained by the ongoing editing and selection happening at day 4, and a more stable reduction by days 10 and 17 (Fig. 2d).

As an orthogonal, DNA-level check that PTC-induced knockdown produces real fitness consequences rather than being a transcript-level artifact, we correlated each PTC guide’s bulk gDNA representation (log2 fold-change vs. the plasmid library) against its target gene’s DepMap essentiality score. This correlation is essentially absent at day 4 (r = 0.07, p = 0.14) but strengthens steadily thereafter (r = 0.17, p = 2.2 × 10⁻⁴ at day 10; r = 0.28, p = 5.5 × 10⁻¹⁰ at day 17, n = 475 designs), consistent with cells carrying PTC guides against more essential genes progressively dropping out of the pool as NMD-induced loss of function accumulates (Fig. 2e).

Because the Flex probe panel resolves multiple probes per gene, including probes spanning specific exon-exon junctions representative of mature transcripts strongly affected by NMD, we further asked whether PTC-induced transcript loss is distributed evenly across a gene’s probes or concentrated at particular probes. Of the 82 day 10 & 17 combined hits, 67 had sufficient probe-level coverage to test (20 UMIs per probe). A probe-composition test comparing each probe’s share of total gene UMI between PTC and proximal-control cells identified five genes (*CD2BP2*, *ELAVL1*, *GNA11*, *KNOP1*, *TRMT2A*) with a significant, non-uniform shift in probe usage (Fig. 2b,f). In the two of these five hits whose probe set includes an exon-exon junction-spanning probe (*CD2BP2, TRMT2A*), that junction probe shows the largest depletion among the gene’s own probes, consistent with disproportionate loss of correctly spliced, mature transcript; across the full panel of 190 probe-level tests this trend is directionally consistent but does not reach genome-wide significance (Mann–Whitney p = 0.31), so we interpret it as a feature of these strongest hits rather than a universal signature of NMD. Differential probe usage for *ELAVL1*, *GNA11* and *KNOP1* however may be stochastic in nature, a feature that has previously been highlighted as a limitation to Flex-based read-outs^26^.

For genes with guides targeting multiple positions (102 of the 225 target genes), whether a given PTC triggers detectable knockdown is strongly pegRNA-dependent: 58 of these 102 genes show no significant guide at all, and of the 44 with at least one hit, three-quarters (33/44) have exactly one significant position among their tested designs, with 9 genes significant at all tested positions (Supplementary Fig. 2a). Regardless of guide quality, this finding is consistent with the classical exon-junction-complex model of nonsense-mediated decay, significance is weakly but measurably associated with a PTC’s distance from the start codon (Spearman ρ = −0.11, p = 0.036, n = 356; Supplementary Fig. 2b), while there is no detectable relationship with distance to the final exon–exon junction (ρ = −0.02, p = 0.74; Supplementary Fig. 2c).

These results show that PEP-seq sensitively and reproducibly detects PTC-induced NMD, while revealing substantial guide-to-guide variation in the strength of the resulting transcript-level phenotype.

### Fitness and gene expression changes induced by eQTLs (Figure 3)

By leveraging paired genetic and transcriptomic datasets, statistical genetics can infer relationships between genetic variants and gene expression, identifying expression quantitative trait loci (eQTLs). However, these associations do not establish causality or resolve the directly affected gene(s), making prime editing a particularly well-suited approach to test candidate alleles cis-regulatory effects.

**Figure 3.**
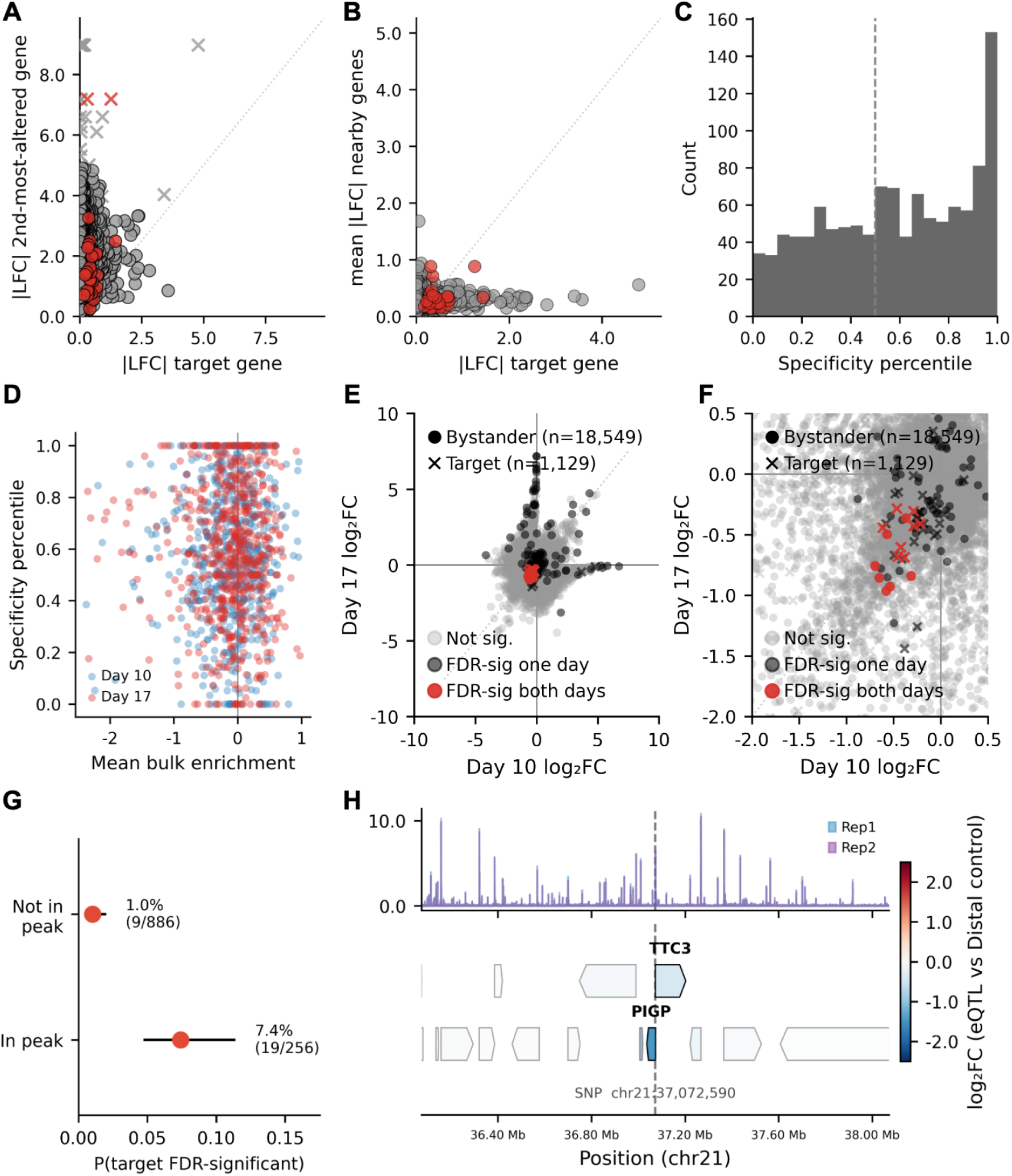
Fitness and gene expression changes induced by eQTLs **(a)** cis-eQTL target-gene specificity, day 17: for 1,146 guide × target-gene pairs (601 guides), each target gene’s |log2FC| plotted against the |log2FC| of the single most-altered non-target ("bystander") gene in the same ±1 Mb window, coloured by target FDR significance. **(b)** The same 1,146 pairs, plotting target |log2FC| against the mean |log2FC| across all bystander genes in the window. **(c)** Distribution of the locus-size-corrected specificity percentile (Methods) at day 17 (n = 1,146). **(d)** cis-eQTL specificity plotted against bulk enrichment or depletion (mean of scRNA-seq and gDNA representation signals), at day 10 and day 17. **(e)** Day 10 vs. day 17 reproducibility of cis-eQTL locus-wide effects, for 19,678 (guide, gene) pairs resolvable at both timepoints (644 guides, 5,286 genes), distinguishing target genes from bystanders and coloured by a three-tier FDR scheme. **(f)** Zoomed view of (e), restricted to the negative-log2FC region. **(g)** Proportion of target genes FDR-significant at day 17, by whether the eQTL SNP sits in a called ATAC peak (present in both replicates) or not (n = 1,142 pairs), with Wilson 95% confidence intervals. The day 10 result is reported in the text. **(h)** Representative eQTL locus on chromosome 21 (chr21:37,072,590 A>G): bulk HEK293T-PE7 ATAC-seq signal (two replicates) across the window, with gene models below coloured by log2 fold-change (eQTL guide vs. distal synonymous control, day 17). The variant sits within a reproducible ATAC peak positioned between the transcription start sites of the locus’s two annotated target genes, *PIGP* and *TTC3*.

Prioritising variants from the EMBL eQTL catalogue, we nominated 839 cis-eQTLs (with <1Mb between the variant and gene target) purportedly linked to 701 unique genes (using up to 3 pegRNA designs per edit). Of the 386 distinct genomic positions, 52.1% were inferred to affect multiple genes.

For each eQTL guide, we tested every coordinate-resolved gene within ±1 Mb of the edit site against a distal synonymous-control background, allowing direct comparison of the guide’s intended target(s) gene against every other gene sharing its locus (that we refer to as "bystander" genes). On a purely raw-effect-size basis, suspected target genes are usually not the most strongly perturbed gene in their own window: at day 17, only 2.4% of 1,146 tested guide–target pairs reach statistical significance for the target, and 97.8% of pairs show a larger absolute log2 fold-change in some bystander gene than in the target itself (Fig. 3a). However, even where a bystander’s raw effect size exceeds the target’s, the mean effect size across all bystanders in the window remains small, and significant targets cluster near this low-bystander-effect region (Fig. 3b). Day 10 results are quantitatively very similar (Supplementary Fig. 3a,b). In short, a variant’s target gene is rarely the largest raw effect in its locus, but is often the most statistically confident one.

To account for this, we developed a locus-size-corrected specificity metric: for each target gene, the fraction of its locus’s own bystander genes that it outranks in estimation precision (a signed Wald z-score derived from each gene’s own fitted log2 fold-change and p-value), computed independently within each guide’s own window and day so that guides at loci with different numbers of nearby genes remain directly comparable. By this measure, a majority of target genes exceed the 50% (uniform-null) expectation at both day 17 (median specificity percentile 0.60; 59.0% of pairs above 0.5) and day 10 (median 0.59; 57.9% above 0.5), with no significant difference between the two days (two-sample Kolmogorov–Smirnov D = 0.055, p = 0.056; Fig. 3c, Supplementary Fig. 3c). This specificity metric is independent of a guide’s gDNA enrichment at either timepoint (day 10: r = 0.02, p = 0.63, n = 605; day 17: r = −0.05, p = 0.23, n = 601; Fig. 3d), arguing that recovered target specificity reflects genuine regulatory signal rather than a proxy for cell-fitness or growth effects.

Because up to three pegRNAs were designed for a single edit position, we also asked whether a target gene’s significance call is reproduced across its guide replicates (Supplementary Fig. 3d,e). When all three guides were tested, the majority show no significant replicate at all for the target gene (77–85%, day 10/17), indicating that a target’s significance call is often guide-specific rather than a robust property of the edited position. Bystander genes replicate even less often (∼99% zero-replicate), consistent with most bystander hits reflecting statistical noise rather than a reproducible effect.

Restricting to genes with a resolvable fit at both timepoints (19,678 guide–gene pairs across 644 eQTL guides and 5,286 genes), effects that replicate across days are rare but highly directionally consistent: only 15 pairs (8 target genes, 7 bystanders) reach significance at both day 10 and day 17, and all 15 (100%) are downregulated on both days (Fig. 3e,f), indicating that genuine, twice-replicated cis-regulatory effects in this dataset are consistently repressive. A matched trans background (30 random genes/guide, ≥1 Mb from the edit site or on a different chromosome) shows this reproducibility is largely cis-specific: only 1 of 17,770 pairs replicate (Supplementary Fig. 3f).

Finally, we asked whether local chromatin state at the variant itself predicts whether its target gene reaches significance, using bulk ATAC-seq performed on unedited HEK293T-PE7 cells. As continuous ATAC pileup signal is dominated by counting noise at the depths achieved here (Methods), we classified each variant as sitting inside or outside a reproducibly called ATAC peak (present in both biological replicates). This binary accessibility state predicts whether the target gene reaches significance: at day 17, target genes are impacted 7.4% of the time when the variant sits in a peak versus 1.0% when it does not (odds ratio = 7.8; cluster-robust p = 3.1 × 10⁻⁵) (Fig. 3g), with a directionally consistent but smaller effect at day 10 (2.7% versus 1.0%; odds ratio = 2.7; p = 0.055). Figure 3h shows a representative example: an eQTL on chromosome 21 (chr21:37,072,590 A>G) whose variant sits within a sharp ATAC-seq peak present in both replicates, positioned less than 1 kb from the transcription start sites of its two annotated target genes: 481 bp from the transcription start site of *PIGP*, and 664 bp from the transcription start site of *TTC3*. At day 17, both target genes are significantly downregulated: *PIGP* strongly (log2 fold-change = −1.44, q = 1.1 × 10⁻¹⁰) and *TTC3* more modestly but still significantly (log2 fold-change = −0.41, q = 3.0 × 10⁻³), illustrating that a single accessible variant can simultaneously perturb both genes flanking a shared regulatory region.

These results show that reproducible eQTL effects tested via PEP-seq are sparse but locally specific, enriched at accessible regulatory elements, and can extend to multiple genes within the same cis-regulatory neighbourhood.

## Discussion

This study establishes a framework for pooled prime editing with single-cell readouts. This assay can be scaled to thousands of independent endogenous variants while maintaining the specificity and statistical power needed to distinguish intended on-target effects from locus-wide bystander variation. The platform recovers an expected, well-characterized phenotype (PTC-induced NMD) with high sensitivity and reproducibility across independent timepoints. In this context, we hypothesize that our platform could also be used in whole-genome knock-out experiments, without the experimental variability introduced by CRISPR non-homologous end-joining^27^. Engineered prime edited knock-outs, provided they can be inserted with high efficiency, could hereby further reduce the cellular coverage required to study gene regulatory networks with confidence.

Our platform extends naturally to a more challenging class of variant: candidate cis-eQTLs where effect sizes are smaller and bystander effects are common. We further introduced a locus-size-corrected specificity metric to assess variant effect and demonstrate that local chromatin accessibility predicts detection probability. Furthermore, the genes that were significant across both later time points showed robust downregulation and were often the target genes of eQTLs. We hypothesise that as cell lines stabilize their gene expression patterns over time in response to the editing process, the robust effects we observe are genuine eQTLs, whereas non-eQTL differentially expressed genes (DEGs) are transient effects originating from media changes, environment, cell state, and stochastic fluctuations. Although, to support this, a much larger study across multiple cell types and contexts should be carried out. A small exploratory panel of candidate splicing-QTL variants was further included in the library design to test for probe composition-level signal orthogonal to whole-gene expression (Methods), which did not expose any hits. This finding underlines the importance of exploring this variant class with single-cell technologies that can resolve full-length transcripts, either via library preparation^28^ or long-read sequencing^29^. In parallel, multi-omic profiling of open chromatin and the transcriptome could further advance non-coding variant screens^30^. Such an approach would resolve the regulatory state of each cell and allow variants to be called directly from accessible regions, making it possible to regress out confounding effects of heterogeneous editing^31^.

Our study also highlights several practical considerations for future variant-to-function screens. Independent pegRNAs targeting the same variant did not always reproduce the same eQTL signature, consistent with variation in pegRNA activity or editing efficiency, supporting the use of multiple pegRNAs per variant. Temporal sampling further suggested that day 4 was too early to capture stable regulatory effects, whereas days 10 and 17 produced broadly concordant signatures. For PEP-seq experiments, we therefore favour concentrating sequencing depth at a single, sufficiently late timepoint, equating to day 10 in our system, while increasing both the number of cells and independent pegRNAs assayed per variant. These parameters will nevertheless depend on cell type, editing kinetics and expected effect size, with stronger perturbations likely requiring substantially fewer cells for confident detection.

A core limitation of variant-to-function screens is the enormous number of candidate variants that must be tested genome-wide, substantially increasing experimental complexity and cost. Compressed screening strategies may provide an alternative, allowing multiple perturbations to be assayed per cell and computationally resolving their individual effects by exploiting the sparsity of biological responses^32^. Because association signals typically implicate many correlated variants, only a subset of variants within GWAS and eQTL loci are expected to be causal^33,34^, suggesting that appropriately calibrated high-MOI designs could similarly enable efficient recovery of sparse variant effects. Sequencing requirements may be reduced further by restricting transcriptomic measurements to informative gene sets, an approach shown to decrease sequencing depth by up to ∼50-fold in targeted Perturb-seq^35^. Probe-based platforms such as GEM-X Flex v2 and VIPerturb-seq provide a natural framework for such targeted measurements through customizable probe-based capture^15,36^.

Finally, with the recent popularity of DNA foundation models^37–39^, whereby paired genetic and epigenetic data is utilized to learn the relationship between variant and downstream regulatory consequence, we expect techniques like PEP-seq to become increasingly important. Such DNA foundation models typically rely on population scale biobank resources, which are limited by genetic variation present in the study group^40,41^. Prime editing perturb-seq offers an alternative: providing the cell model in question is believed to be representative enough of the tissue of interest, one can then directly test any variant of interest prospectively — even variants that have never been observed^42^.

## Methods

### Key resources and materials

Oligonucleotide sequences can be found in Supplementary Table 1, the list of reagents can be found in Supplementary Table 2 and the CROP-seq vector sequence can be found in Supplementary File 1.

### HEK293T sub-culture routine

HEK293T cells were cultured in DMEM+GlutaMAX supplemented with 10% fetal bovine serum (FBS) and 1x antibiotic-antimycotic at 37℃ in a humidified incubator supplemented with 5% CO_2_. Cells were passaged every 2-4 days using TripLE or Trypsin, following the manufacturer’s instructions. Cell counting was performed on a Luna FX7 Automated cell counter with acridine orange/propidium iodide (AO/PI) staining.

### HEK293T-PE7 cell line generation

HEK293T cells were transduced with EF1a-PE7-2A-blasticidin and EF1a-PE7-2A-GFP lentiviruses (Supplementary Files 1 and 2, cloned and ordered via VectorBuilder), each with a MOI of 2 in the presence of 8 µg/mL polybrene. The next day, cells were expanded following sub-culture passaging and selected for with 5 µg/mL of blasticidin S. After a week of selection, GFP^+^cells were sorted using a Bigfoot Spectral Cell Sorter with a 100 µm nozzle. To select for cells with editing activity, GFP^+^ cells were nucleofected with 8µg of pegRNA programming a single nucleotide variant catalysing the transition from GFP to BFP (Supplementary Table 1) using a SF Cell Line 4D-Nucleofector LV Kit L in a Lonza 4D-nucleofector with programme CM-130. BFP^+^ cells were selected on the Bigfoot Spectral Cell Sorter and individual clones were derived using limited dilutions. Each clone was further nucleofected as described above but using a pegRNA catalysing the transition from BFP back to GFP (Supplementary Table 1), and the proportion of GFP^+^ cells in each clonal population was calculated by Flow cytometry (Bigfoot Spectral Cell Sorter). The clone (HEK293T-PE7) with highest editing efficiency was further expanded and banked for further prime editing assays.

### Transduction and cell fixing

Lentiviral pegRNA libraries were manufactured using VectorBuilder. Four million HEK293T-PE7 cells were seeded in each of four separate T75 flasks. One seed culture was used to estimate cell numbers and calculate the lentiviral amount required for transduction. 24 hours after seeding, the remaining three flasks were transduced with the lentiviral pegRNA library with an MOI of 0.1. Starting 24 hours after transduction, and throughout the rest of the experiment, transduced cells were selected by supplementing the media with 1µg/ml of Puromycin Dihydrochloride. At each of the harvesting timepoints (3, 10 and 17-days post-transduction), for each replicate at least 2 million cells were kept in culture, 1-2 million cells were stored at -80°C for posterior gDNA extraction, and 1-2 million cells were fixed and processed for 10X Flex Apex with GEM-X Flex Sample Preparation v2 Kit, following the manufacturer’s instructions (10X Genomics, CG000782 protocol). All samples were fixed for 18 hours and 40 minutes and stored at -80°C.

### Single-cell sequencing of prime-edited cells

To assay edited cells, the 10x Flex Apex protocol (CG000834, RevB) was carried out with the VIPerturb-seq implementation for guide capture, as described below. LHS and RHS VIPerturb-seq probes were ordered as Oligo Pools (Integrated DNA Technologies) and spiked into the Probe Hybridization Mix following recommendations from the manufacturer. Each sample was split into 10 reactions for sample barcode hybridization, after which all samples were pooled prior to washing using the pooled wash workflow. For encapsulation, cells were loaded across four GEM-X wells (three wells loaded with 600,000 cells, and one well loaded with 180,000 cells). During library preparation, the pre-amplification reaction was supplemented with 1 µl of 10 µM GDO primer. Eight indexing PCRs were performed for each GEM-X well, four of them amplifying the gene expression library (GEX) and four of them amplifying the guide capture libraries (GDO), as per the VIPerturb-seq protocol. Libraries were pooled with a GDO to GEX molarity ratio of 1:10 and sequenced on a P4 flow cell (for sequencing the single 180,000-cell GEM-X well) and a 25B flow cell (for sequencing of the remaining 600,000-cell GEM-X wells) with 2% PhiX spiked-in and the following run configuration: Read 1: 54 cycles; Index 1 (i7): 10 cycles; Index 2 (i5): 0 cycles; Read 2: 50 cycles.

### Genomic DNA harvesting and lentiviral insert sequencing

The genomic DNA and RNA were harvested from frozen pellets using the Qiagen Allprep DNA/RNA kit, and quantified on Qubit DNA and RNA high-sensitivity kits. The integrated lentiviral construct containing the pegRNA sequence and VIPerturb-seq barcodes was then amplified from up to 1.25 µg of DNA per 100 µl PCR reaction using the NEBNext Ultra II PCR mastermix supplemented with 2 mM of MgCl_2_ and 0.6 µM forward and reverse primers (Supplementary Table 1). The product was amplified for 16 cycles, using an annealing temperature of 72°C, and each reaction was pooled for each sample and 100 µl were purified at a 1:1 volumetric ratio using SPRIselect and eluted in 30 µl nuclease-free water. For the library amplification, 1 µl of glycerol stock was added as an input and amplified in 50 µl NEBNext Ultra II PCR reaction, amplified for 8 cycles (annealing temperature of 68°C) and cleaned-up with the same volumetric ratio as the other libraries. A subsequent indexing PCR (Supplementary Table 1) was performed by amplifying 1 µl of product in a 50 µl NEBNext Ultra II reaction for 10 cycles, purified using a 1:1 volumetric ratio of SPRIselect beads. The libraries were then pooled equimolar and sequenced on a P2 300-cycle flow-cell with 30% PhiX spiked-in.

### Bulk ATAC-seq library preparation

Bulk ATAC-seq was performed on unedited HEK293T-PE7 cells in two biological replicates using the Diagenode Bulk ATAC-seq kit, following the manufacturer’s workflow with digitonin-based nuclei isolation. Approximately 50,000-60,000 nuclei were processed per replicate; cells were lysed in ATAC Lysis Buffer supplemented with digitonin, washed, and nuclei were immediately subjected to tagmentation. Tagmentation was performed for 30 min at 37°C, after which DNA was purified and libraries were indexed and PCR amplified, with the number of additional amplification cycles determined by qPCR; four additional cycles were used, giving nine total amplification cycles. Libraries were size-selected using AMPure XP beads and assessed for concentration and fragment-size distribution prior to sequencing. Libraries were sequenced on an Illumina NextSeq 1000/2000 using a P2 100-cycle kit with 2 × 50-bp paired-end reads and dual 8-bp indexes.

### eQTL variant prioritisation

Candidate eQTLs were sourced from the EMBL eQTL catalogue, applying filters for significant associations (P < 0.001) and substantial effect sizes (|β| > 1). Variant-gene pairs across datasets were consolidated, retaining only those with concordant effect directions. Annotation was performed using HEK293T gene expression data and coverage by 10x Genomics Flex probes. Variants were prioritised via a composite metric based on median standardized effect size (β/SE), HEK293T expression levels, replication across cohorts, fine-mapping posterior inclusion probability (PIP), credible-set dimensions, LD profile, and statistical significance.

Selection criteria required observation in at least 4 datasets, targeting of Flex-compatible genes expressed in >30% of HEK293T cells (mean expression >0.2), a peak PIP ≥0.7, and a |median Z| ≥5. Candidates meeting a composite score threshold of 2 were then ranked for final inclusion.

### sQTL identification and reclassification

The sQTL variants were not the product of an independent sQTL selection pipeline: every sQTL locus is a member of the same pool of 557 candidate cis-regulatory variants selected under the eQTL variant-prioritisation criteria above, and was originally designed and synthesised as an eQTL pegRNA. After variant selection, target-gene assignment for all 557 variants was re-audited against the EMBL eQTL Catalogue’s reported quantification method for each dataset, classifying evidence into three tiers: expression (gene expression, microarray), splicing (exon, transcript, transcript-ratio, and intron-excision/leafcutter quantifications), and protein (aptamer-based); and re-applying the original significance filter (|β| > 1, P < 1×10⁻³) within the best-available tier per variant, in priority order expression > splicing > protein. For 171 of the 557 variants (30.7%), no expression-tier association survived this filter but a splicing-tier association did; these variants were reclassified from eQTL to sQTL, and their target gene(s) reassigned to the best-supported gene(s) under the splicing evidence tier (a single gene for 140 variants; two to four co-regulated genes for the remaining 31). The pegRNAs already designed for these loci under the eQTL category’s own efficiency and length thresholds (see “Final variant and pegRNA selection” below) were re-categorised as sQTL (348 pegRNAs across the 171 loci: 65 loci with one pegRNA, 35 with two, and 71 with three).

### Selection of genes for PTC and synonymous variants

Genes for targeted nonsense-mediated decay induction were selected based on perturbation profiles in HEK293T cells from the X-Atlas/Orion dataset. A global impact score was derived from the e-distance in PCA space between gene-specific perturbations and non-targeting controls. Candidates were annotated with DepMap common-essential designations^43^ and filtered for inclusion in the 10x Flex probe set. The final 400-gene panel comprised the 200 highest-impact DepMap essential genes and the 200 highest-impact non-essential genes, ensuring a mutually exclusive set for downstream analysis.

### PTC and synonymous control variant design

Transcript selection prioritised the MANE Select transcript^44^, followed by Ensembl canonical or the longest CDS. Premature termination codon (PTC) variants were restricted to sites likely to trigger nonsense-mediated decay, defined as >150 nt from the CDS start and >55 nt upstream of the last exon junction. For each gene, up to four PTCs were designed, specifically targeting third-position tyrosine codon (TAT/TAC) transitions to TAA; single-nucleotide nonsense substitutions were utilised where tyrosine targets were unavailable. Stop codons followed by a+4 cytosine were excluded. Synonymous controls were matched to the same nucleotide position (TAT↔TAC).

### Final variant and pegRNA selection

PegRNA designs were generated using PRIDICT2 and filtered for predicted HEK293T editing efficiencies ≥70 and lengths ≤150 nt. For PTC/control variants, only sites with viable designs for both variant types were selected. These were ranked by the predicted PTC editing efficiency, selecting up to three positions per gene with non-overlapping spacer sequences. Synonymous controls were designed to share spacers with their paired PTC whenever possible. For eQTL variants, up to three pegRNAs with distinct spacers were selected per site, ranked by predicted editing scores in HEK293T cells and adhering to the same efficiency and length thresholds.

### Library oligonucleotide sequence assembly

Selected pegRNAs were then converted into an oligonucleotide, adding flanking homology arms for PCR amplification and Gibson assembly^45^ (“GGAAAGGACGAAACACC” at the 5’ end and “AAGCTTGGCGGGATCCG” at the 3’ end) into the digested pLV-EF1APuro-WPRE-3LTR_hU6BsmBI-stuffer-BsmBI_3LTR backbone synthesized by VectorBuilder. Following the 5’ adapter, the pegRNA sequence generated by PRIDICT2 was appended, followed by a poly(T) transcription termination sequence “TTTTTTT” and a variable-length stuffer to normalise oligonucleotide length to 250bp “GCTATCACCTCTAAAGCCGCACAGTAAGCGTACTAGCCCTCATACTAATGTTAGGCTGCC”. A unique combination of RHS and LHS barcodes was then appended, 3’ of the stuffer. Lentivirus production and quality control was performed by VectorBuilder.

### Single-cell data pre-processing with Cell Ranger

Fastq files were obtained using bcl2fastq and were processed using Cell Ranger v10.1.0 (cellranger multi). Gene-expression libraries were analysed using the Flex-v2-R1 chemistry and the Chromium Human Transcriptome Probe Set v2.0.0 (GRCh38-2024-A). CRISPR guide-capture libraries were processed using a custom feature reference with min-crispr-umi=1; twelve gene-expression and twelve corresponding guide-capture FASTQ sets, four per GEM-X well, were supplied and samples were demultiplexed according to their specified probe-barcode combinations.

### Guide calling and guide multiplicity estimation

Guide (protospacer) identity was assigned per cell by Cell Ranger v10.1.0’s crispr_analysis pipeline (min-crispr-umi=1) from the guide-capture library, yielding a per-cell record of the number of distinct guide features detected and their associated UMI counts; no additional custom UMI threshold was applied on top of Cell Ranger’s own call. A guide multiplicity (GM) count is defined directly as a cell with zero called guides (GM = 0) is retained in the object but excluded from every downstream guide-level test, and a cell with more than one called guide (GM > 1) carries a combined guide identity. Guide identifiers were corrected for two known label issues before any downstream use: (i) a subset of variants originally labelled eQTL were relabelled sQTL (and their target-gene identity revised) based on the independent re-audit described above (sQTL identification and reclassification); and (ii) guide-replicate suffixes were renormalised to a stable, alphabetically sorted g1/g2/g3 scheme per genomic position. Each guide’s target-gene set was resolved by mapping its Ensembl gene ID(s) to HGNC symbols via the Cell Ranger gene-expression feature reference. Nearby genes for every guide’s edit position were pre-computed as a ±1 Mb window lookup built from the GRCh38 GENCODE reference (joined on Ensembl gene ID, not gene-symbol string, to avoid symbol-vintage mismatches), and this same window is reused for every proximal/distal, bystander, and locus-based analysis described below.

### Single-cell QC filtering and normalization

Gene-expression and CRISPR guide-capture count matrices were loaded per (lane, sample) from Cell Ranger’s filtered feature-barcode matrix outputs, concatenated across all lanes and samples, and standard QC metrics (genes detected per cell, total UMI counts, mitochondrial read percentage) were computed. Cells were retained if total UMI count fell in between 5,000 and 75,000 and mitochondrial fraction was ≤5%, reducing 1,200,369 cells to 871,203 cells (72.6%) across all three timepoints. Following filtering, raw counts were preserved for all downstream negative-binomial testing, and a parallel log-normalized representation (normalization to 10,000 counts/cell followed by log1p transformation) was used only for visualization and clustering: 2,000 highly variable genes were selected (blocked by sample), reduced to 30 principal components, and embedded using a standard neighbor-graph/UMAP pipeline (10 principal components). A second, independent guide-level QC layer flags cells carrying any pegRNA design whose day-0 plasmid-library integrity fell below adopted thresholds (minimum supporting reads, barcode/spacer concordance, and full-construct sequencing-fidelity thresholds, tuned by elbow-diagnostic against a less stringent default); this flag identifies 122,093 of 871,203 QC-passing cells (14.0%), fairly evenly distributed across timepoints (10.3% / 15.9% / 15.6% at days 4/10/17) leaving 749,110 cells for final downstream analysis.

### Guide-induced cluster-occupancy test

Leiden clustering (scanpy, flavor="igraph", resolution = 1) was run on the post-QC cell atlas; recomputing after GM = 1 restriction, or with/without low quality guide transduced cells did not change assignments and was not used. For each timepoint group (day 10, day 17, combined) and guide with ≥25 GM = 1 cells, its per-cluster profile was compared to the pooled population via JSD (base-2, bounded [0,1]) against a matched-n null: 200 synthetic multinomial draws of the guide’s own cell count from the pooled cluster proportions. A guide was flagged if its JSD exceeded the 95th percentile of a null self-calibrated from only the subgroup being tested – exact under the global null regardless of within-subgroup cell-count heterogeneity, not approximate. Fitness effect was the guide’s day-17 gDNA log2FC vs. plasmid library (Methods, "gDNA library integrity analyses"); |log2FC| > 1 defined high-fitness-effect.

### Per-cluster enrichment test

Cells from a subgroup’s guides were tallied per cluster and each cluster’s rate compared to the timepoint group’s pooled rate via one-sample binomial test (population rate treated as fixed, given far larger n), BH-corrected across the 18 clusters per timepoint group. Log2 enrichment used a continuity-corrected rate, (k+0.5)/(n+1).

### Negative-binomial differential expression testing

All guide-level differential expression tests (NMD target-gene effects, eQTL cis/bystander effects, and steric distance scans) use a common negative-binomial Wald test, fit per gene on raw integer counts against a binary foreground/background group indicator, with each cell’s total UMI count entered as a fixed library-size offset. Log2 fold-change is the fitted group coefficient divided by ln(2). Two failure modes of this test are handled explicitly: near-separation falls back to a likelihood-ratio test against an intercept-only null, and complete separation (one arm identically zero) is resolved with an exact closed-form test calibrated to avoid the anti-conservative behavior of a boundary-optimum likelihood-ratio test in this regime, with fold-change reported using a Haldane–Anscombe pseudocount anchored to the gene’s pooled observed rate. Gene-level dispersion is pre-estimated once per (dataset, timepoint) from a large distal synonymous-control background pool and held fixed in each individual guide’s test at that timepoint. Multiple-testing correction is Benjamini–Hochberg, applied within each guide’s (or guide × target’s) own family of nearby-gene tests per timepoint (FDR threshold q < 0.05). Day 10 and day 17 are treated as independent test families throughout, except for the NMD analysis (Figure 2), which additionally runs a combined day 10+17 family to maximize power.

### eQTL specificity metric

For each cis-eQTL guide with sufficient variant cells at day 10 and day 17 separately, log2 fold-changes and p-values were computed for every gene within the guide’s own ±1 Mb window against a distal synonymous-control background, using the shared negative-binomial pipeline described above. Within each guide’s own locus and day, p-values across all tested nearby genes were Benjamini–Hochberg-corrected, and each gene’s log2 fold-change/p-value pair was converted to a signed Wald z-score so that ranking is driven by estimation precision rather than raw magnitude alone. For a guide’s declared target gene(s), specificity was scored against the pool of all other (“bystander”) genes tested in the same locus and day as: (1) the ratio of the largest bystander effect size to the target’s own effect size; (2) the mean bystander effect size relative to the target’s own effect; and (3) a locus-size-corrected specificity percentile, defined as the fraction of that locus’s own bystander genes whose precision the target gene’s precision exceeds.

### Bulk ATAC-seq accessibility and peak-overlap analysis

Bulk ATAC-seq reads from the two biological replicates were demultiplexed from BCL files using bcl2fastq. Nextera adapter sequences were removed from both read mates using cutadapt, retaining reads ≥20 bp after trimming. Trimmed paired-end reads were aligned with Bowtie2 using the --very-sensitive preset and a maximum fragment length of 2,000 bp (-X 2000), and alignments were coordinate-sorted and indexed using samtools. PCR duplicates were removed using the samtools fixmate/markdup workflow, comprising name sorting, mate-information correction (fixmate -m), coordinate sorting and duplicate removal (markdup -r). Properly paired alignments with mapping quality ≥30 were retained using samtools view -f 2 -F 1804 -q 30; mitochondrial and chromosome Y alignments were excluded, followed by removal of reads overlapping ENCODE hg38 blacklist v2 regions using bedtools. Peaks were called independently for each biological replicate using MACS3 in paired-end mode (BAMPE). Narrow peaks were called with -q 0.01 --nomodel --call-summits -B --SPMR --keep-dup all, with broad peaks additionally called using --broad; SPMR-normalized pileup tracks generated by MACS3 were retained for quantitative accessibility measurements.

Local chromatin accessibility around eQTL variants was derived from bulk ATAC-seq performed on unedited HEK293T cells (two biological replicates). Continuous local accessibility was summarized as a length-weighted mean pileup value across a fixed window around each position of interest, averaged across replicates. For the SNP-level accessibility-versus-target-effect analysis (Figure 3g), continuous pileup signal was replaced with a binary, reproducibility-gated readout: a variant was classified as sitting within a called ATAC peak only when a peak was independently called in every replicate. This binary accessibility state was compared against target-gene effect size and against FDR significance by weighted least squares and logistic regression respectively, each with two-way cluster-robust standard errors as described above, and group-wise significance rates are additionally summarized with Wilson (rather than normal-approximation) 95% confidence intervals given the small event counts involved.

### Steric and bystander distance-effect analysis

For every guide across all four variant classes with sufficient variant and distal-control cells at a given timepoint, log2 fold-change and significance were computed for every coordinate-resolved gene within the guide’s own ±1 Mb window against its own distal synonymous-control background, using the same pipeline and per-timepoint dispersion described above. Each nearby gene’s signed distance from the edit site was computed gene-body-aware (zero if the edit falls within the gene body, otherwise signed distance to the nearest edge). A guide’s own intended target gene is retained in this pool (flagged accordingly) rather than dropped. Day 10 and day 17 results were pooled for visualization as a density of effect size against genomic distance from the edit site.

### Cluster-robust standard error estimation

The two regressions just described — the ATAC-accessibility analysis above and the distance-effect analysis above — are both fit on (guide, gene) pairs, not independent observations: a given guide contributes one row per nearby gene it is tested against, and a given gene contributes one row per guide tested near it. Treating these rows as independent would understate the true standard errors.

To correct for this, we computed standard errors for both regressions using two-way cluster-robust variance estimation — clustering simultaneously on guide identity and on gene identity, following the Cameron–Gelbach–Miller two-way clustering construction — instead of the model’s default independent-errors covariance. This correction changes only the standard errors and p-values; point estimates are unchanged, and the naive (uncorrected) fit is reported alongside the corrected one in every case.

### Probe count and metadata generation

Probe-level expression matrices were constructed from the Cell Ranger outputs by isolating raw UMI counts from the sample raw probe-barcode matrix files. These matrices were subset to include only high-quality cells identified in the filtered feature matrices. Probe characteristics, comprising gene association, sequence, and splice-junction annotations, were integrated from the probe-set reference. The resulting sparse cell-by-probe matrices were aggregated across the four GEM-X wells, preserving metadata for sample, replicate, timepoint, and well origin.

Probe features were further enriched with genomic coordinates and structural annotation from the Chromium Human Transcriptome Probe Set reference, specifically flagging junction-spanning probes. The final processed dataset was archived as an AnnData object containing raw probe-level UMI counts and comprehensive cell-probe metadata.

### Probe-level composition testing

For target genes with a significant whole-gene expression change (Figure 2b), a complementary test asked whether the change is distributed evenly across a gene’s Flex probes or concentrated at specific probes. For each probe of a tested gene with sufficient combined UMI support, that probe’s share of the gene’s total UMI was compared between perturbed and control cells using a pseudocount-stabilised log-odds test, summarised as a “selective delta”: we calculate the probe’s own usage log2 fold-change minus the mean usage log2 fold-change of its sibling probe. The result is a quantity invariant to uniform, whole-gene decay and sensitive only to genuine shifts in relative probe usage. P-values were pooled per gene (each gene’s minimum probe p-value) and Benjamini–Hochberg-corrected across the tested genes.

### gDNA library integrity analyses

Paired-end sequencing data from the plasmid library and cellular gDNA were retrieved using bcl2fastq and processed using a custom Python pipeline developed in Python 3.10 and utilizing the edlib alignment library (v1.3.9.post1). To identify individual library members, the 50-nt barcode at positions 21-70 of Read 2 was matched, permitting up to four mismatches, while the 20-nt spacer was independently assigned from Read 1 with a two-mismatch tolerance.

Constructs were classified as concordant if the independently called spacer sequence aligned with the barcode-associated expectation; otherwise, they were logged as discordant to assess potential recombination or synthesis errors. The fidelity of the full-length constructs was benchmarked against a reference derived from the 250-nt stitched oligo (TAC + reverse complement). This reference was segmented into distinct functional domains, including the barcode, linker/stuffer/poly-T, PBS, RTT, scaffold, spacer, and constant regions. Each Read 2 sequence was aligned globally using edlib to determine substitution, insertion, and deletion rates. Fidelity was categorized as high (error rate ≤2%) or moderate (error rate ≤8%). Quality control filters excluded plasmid-library members with <20 supporting reads, barcode-spacer concordance <80%, or where high-fidelity reads constituted <50% of the total. This plasmid-level integrity call is separately used to flag single cells carrying a construct with marginal library-level integrity (Methods, “Single-cell QC filtering and normalization”).

## Data and code availability

To be made available upon publication.

## Acknowledgements

We would like to thank Rahul Satija (New York Genome Center) for their help in setting-up VIPerturb-seq and Peter Smibert (10x Genomics) for helpful conversations regarding the implementation of PEP-seq. We would also like to thank Amaia Villas Zornoza, Chantelle Patience and Bianca Ambrosio (Relation Therapeutics) for their assistance with library sequencing; Michal Grochmal and Ron Finn (Relation Therapeutics) for their assistance with data engineering queries; Sorina Maciuca and James Steven Harrison (Relation Therapeutics) for the helpful discussions regarding variant prioritization.

## Contributions

J.D.J. and J.T.K. designed the study. M.A.R., J.D.J. C.J. and J.W.O. performed the experiments. J.D.J. and J.B.C. performed data preprocessing and variant prioritization, and J.T.K. analysed the data. J.D.J., M.A.R., J.W.O. and J.T.K. wrote the manuscript, with input from all authors.

---

**Supplementary Figure 1.**
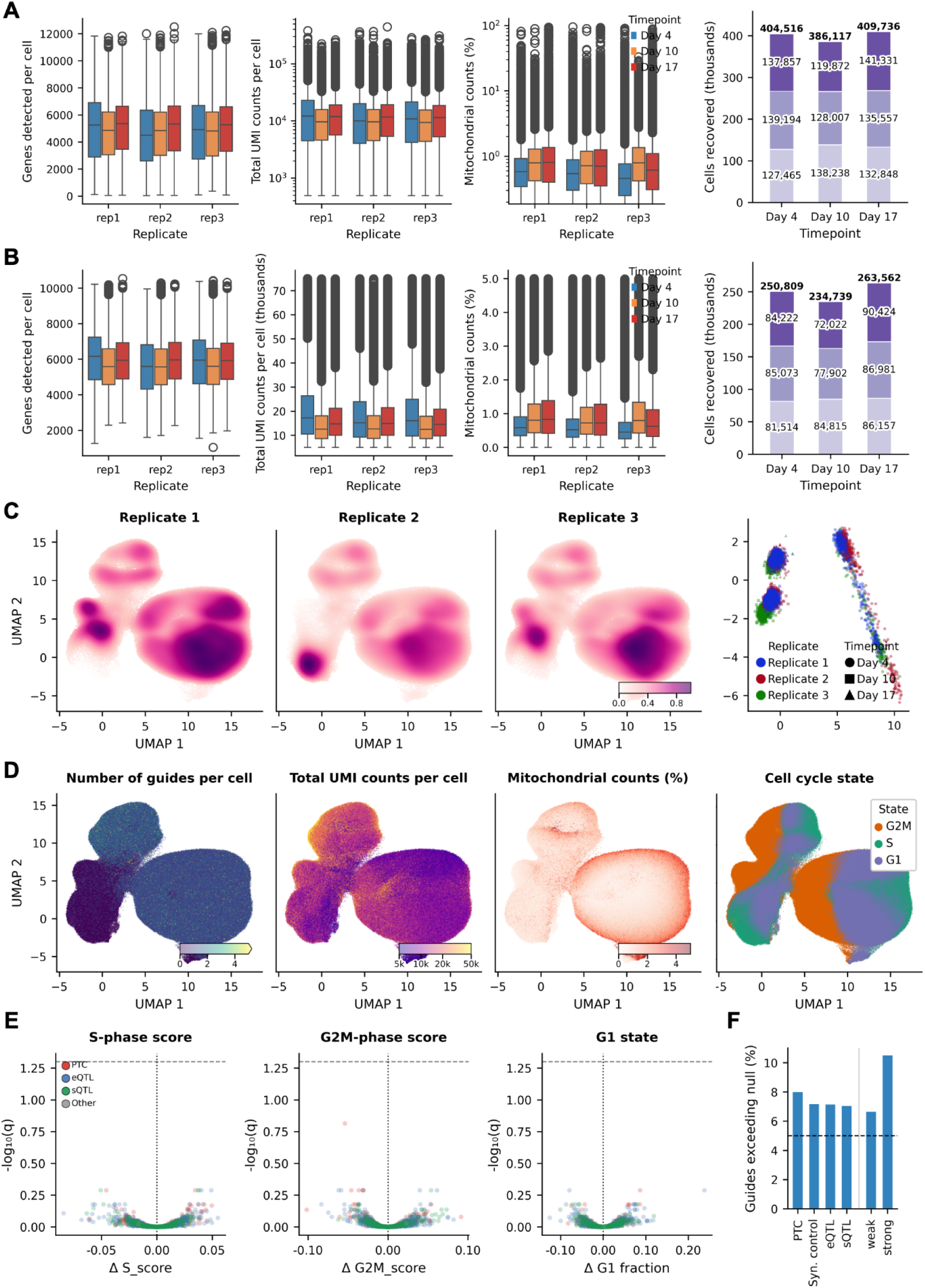
**(S1a)** Cell numbers and single-cell QC metrics for the raw, unfiltered dataset (1,200,369 cells across 9 samples), before the QC threshold used throughout the paper is applied: cells recovered per timepoint and replicate; genes detected, total UMI counts, and mitochondrial percentage per cell; and total UMI counts vs. genes detected, coloured by mitochondrial percentage. **(S1b)** The same panels restricted to 749,110 QC-passing cells that also pass the independent, plasmid-library-based guide-construct-integrity filter (Methods, "Single-cell QC filtering and normalization"). **(S1c)** Batch-effect checks on the same QC-passing population as Figure 1e,f: per-replicate kernel density estimates on the shared UMAP embedding, alongside a pseudobulk PCA of (replicate × guide × timepoint) profiles (GM = 1 cells only). **(S1d)** Assessment of standard QC metrics over UMAP coordinates: number of guides per cell, total UMI counts per cell, mitochondrial counts percentage, and rank-based categorical cell-cycle state (G1/S/G2M) assigned per cell (Methods). **(S1e)** A guide-level test for enrichment or depletion of cell-cycle state (1,643 guides with ≥25 GM = 1 cells, days 10 and 17 pooled). **(S1f)** Cluster-occupancy equidistribution test: for each guide, whether its cells’ occupancy across the 18 Leiden clusters (Fig. 1e) diverges from the population baseline beyond a self-calibrated 95th-percentile null threshold (Methods), summarised as flagged rate by perturbation type and by bulk fitness-effect magnitude (day 10+17 combined).

**Supplementary Figure 2.**
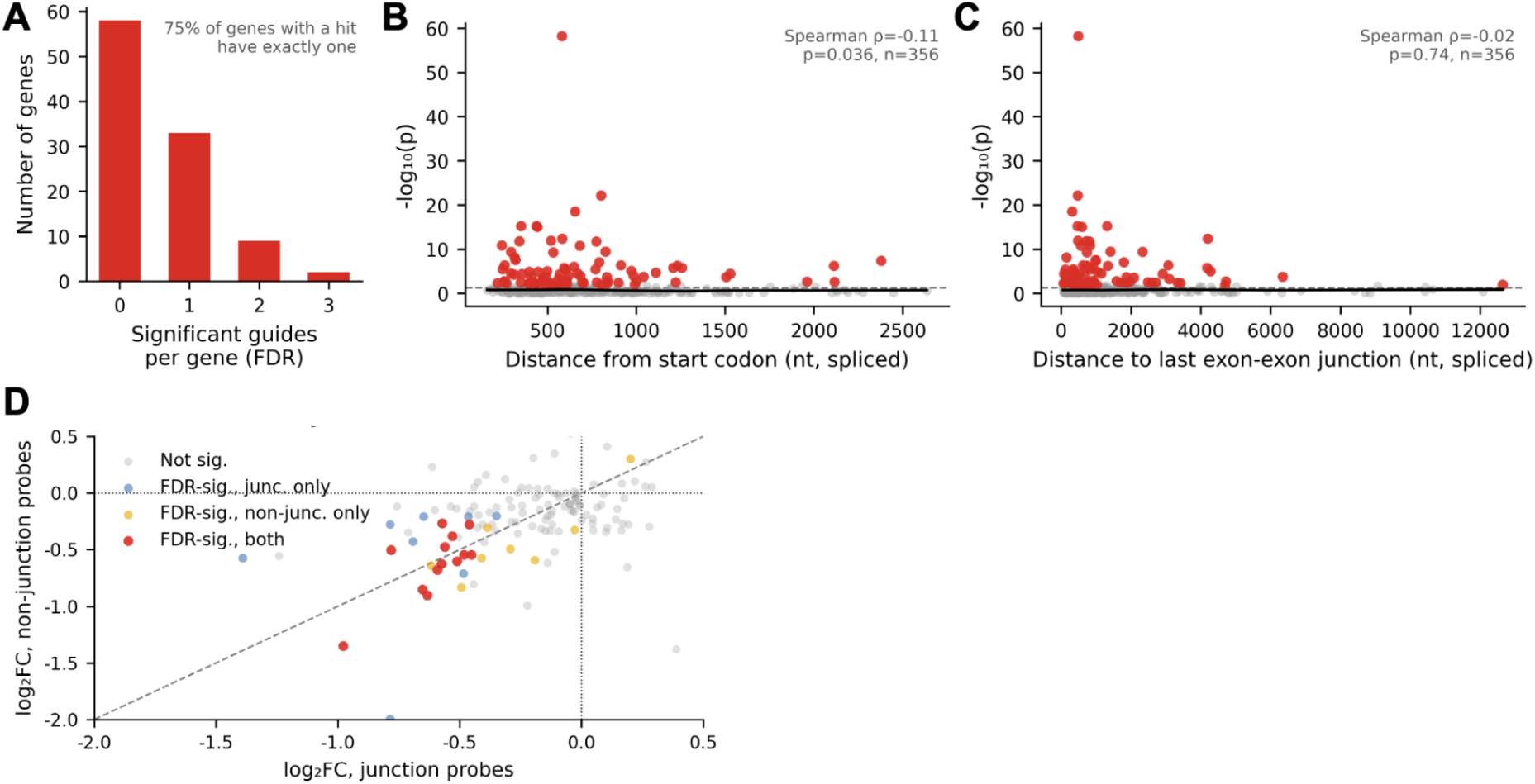
**(S2a)** Guide-level heterogeneity underlying the hit calls in Figure 2b: for the 102 target genes tested at more than one PTC position, the distribution of how many of a gene’s positions were individually FDR-significant. **(S2b)** Relationship between a PTC’s distance from the start codon (in spliced-mRNA nucleotides) and knockdown significance, for the 356 tested positions with a canonical-transcript exon mapping. **(S2c)** Relationship between a PTC’s distance to the final exon–exon junction and knockdown significance, for the same 356 positions. **(S2d)** Junction-vs. non-junction-probe knockdown significance, per PTC guide–target-gene pair (n = 133 pairs, 89 genes): summed junction-spanning-probe log2 fold-change plotted against summed non-junction-probe log2 fold-change (guide vs. proximal synonymous control, day 10+17 combined), coloured by which axis (if either) reaches FDR significance.

**Supplementary Figure 3.**
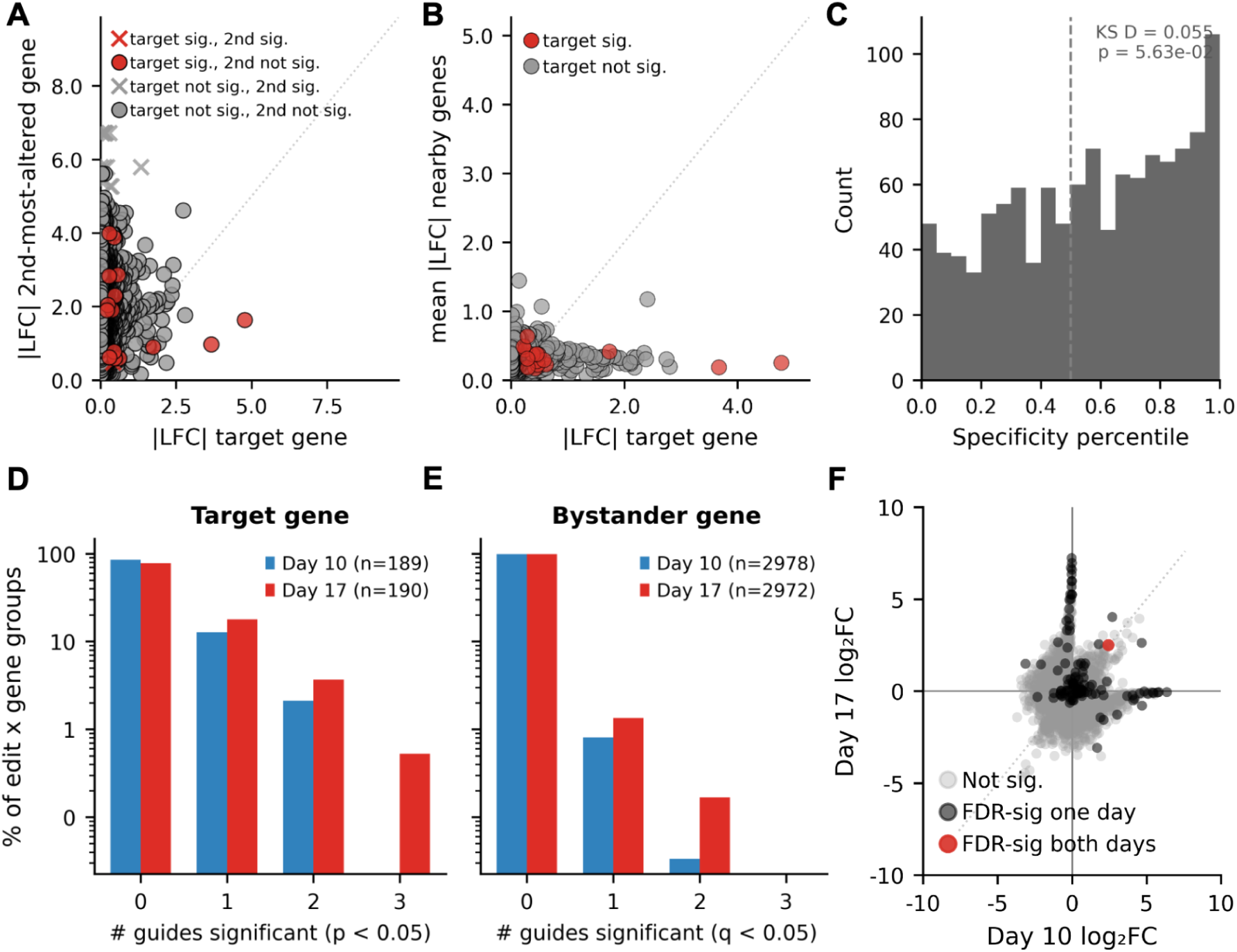
**(S3a–c)** As Figure 3a–c, restricted to day 10 (n = 1,156 guide × target-gene pairs, 605 guides). **(S3d)** Within-edit replication of target-gene significance across independently designed guide replicates (g1–g3): for edit positions with all three tested, the distribution of how many replicates independently reach nominal significance (p < 0.05) for the edit’s own target gene (day 10: n = 189 edit × target-gene groups; day 17: n = 190). **(S3e)** As (S3d), for bystander genes (FDR q < 0.05) at the same edit positions (day 10: n = 2,978 edit × gene groups; day 17: n = 2,972). **(S3f)** Day 10 vs. day 17 concordance for a matched trans background, analogous to Fig. 3e,f but restricted to genes that can never be a guide’s own target (30 randomly sampled genes ≥1 Mb from the edit site or on a different chromosome, per guide, fixed across days; n = 17,770 guide–trans-gene pairs, 644 guides).

**Supplementary Table 1.**
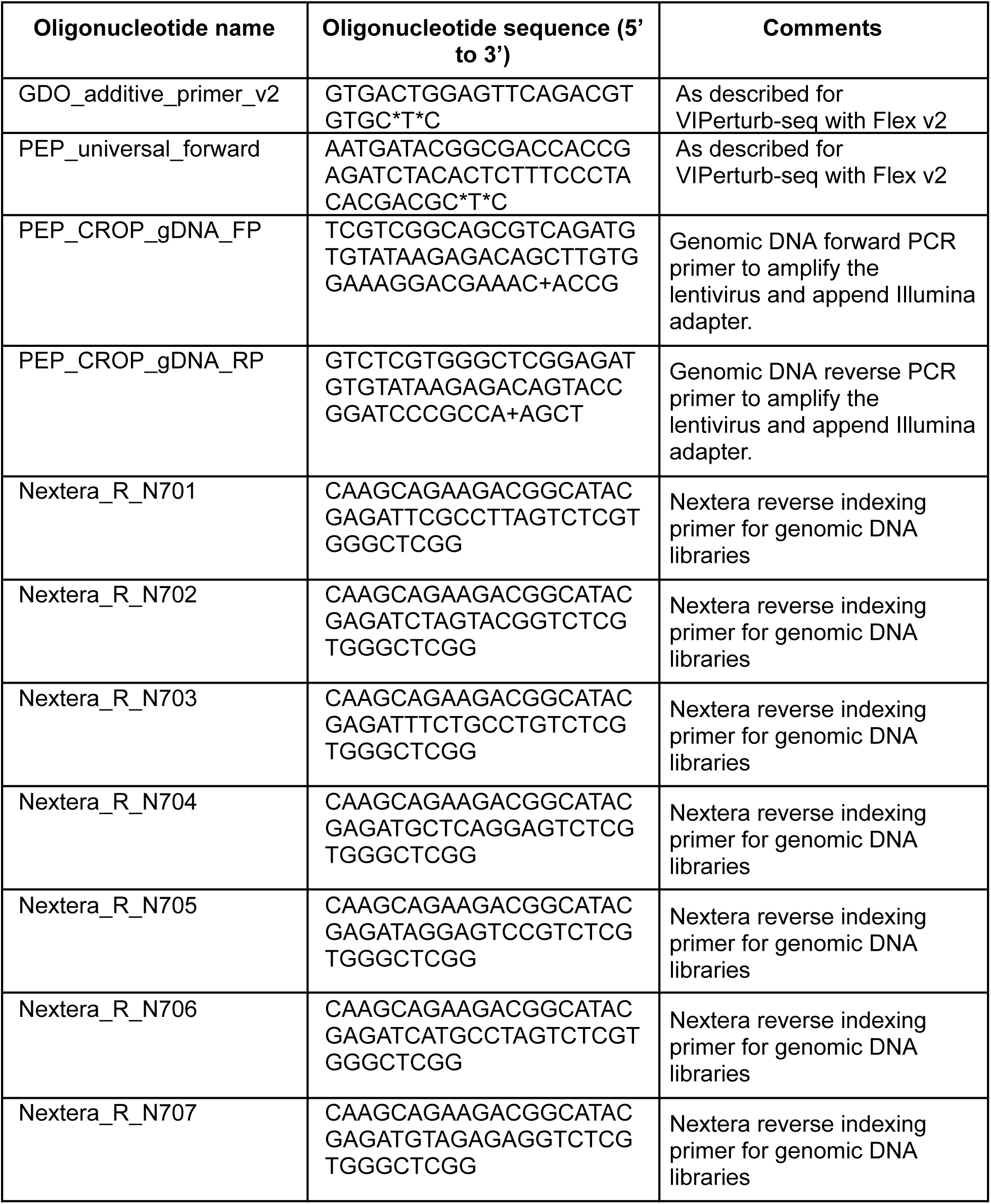
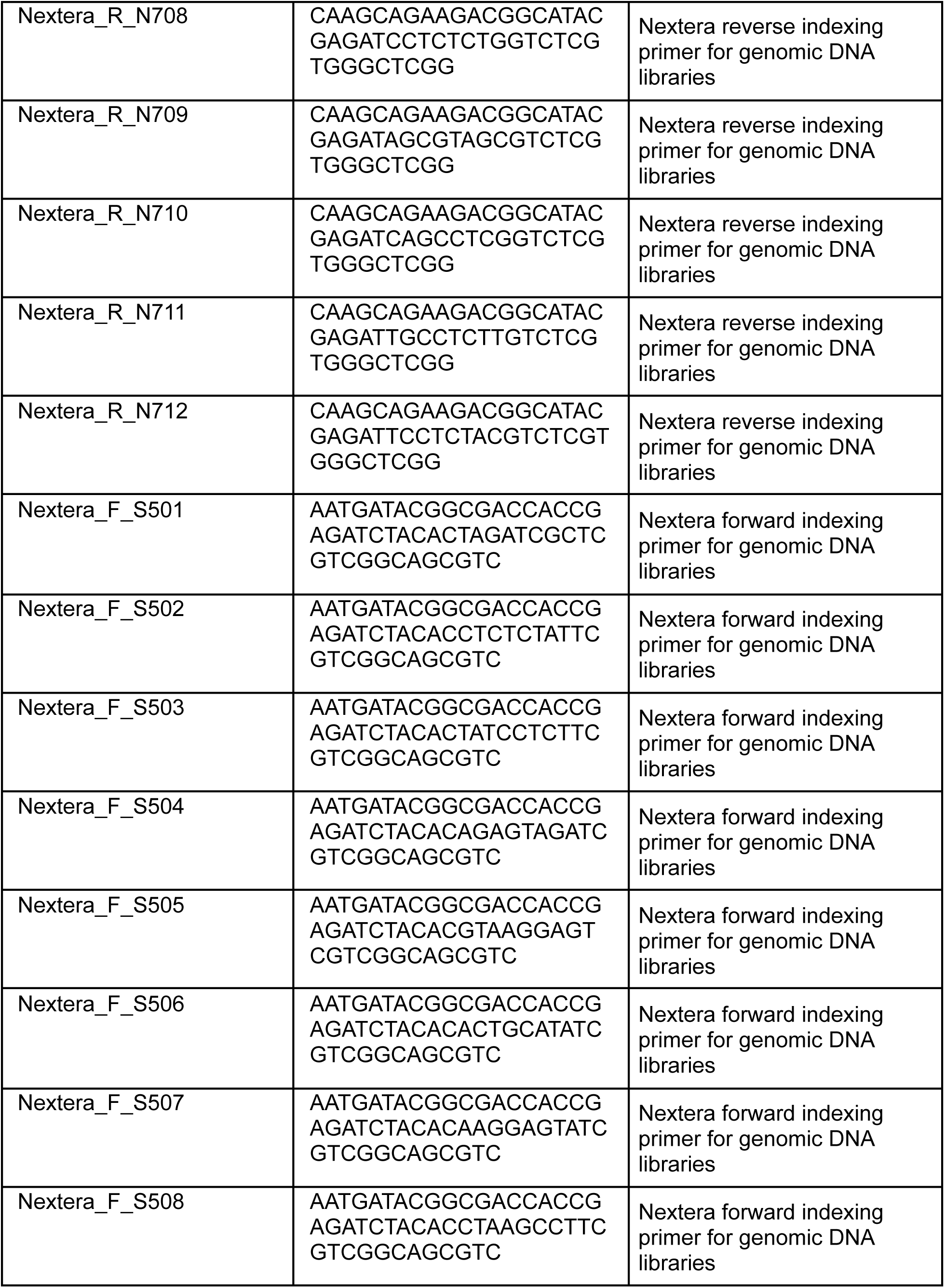
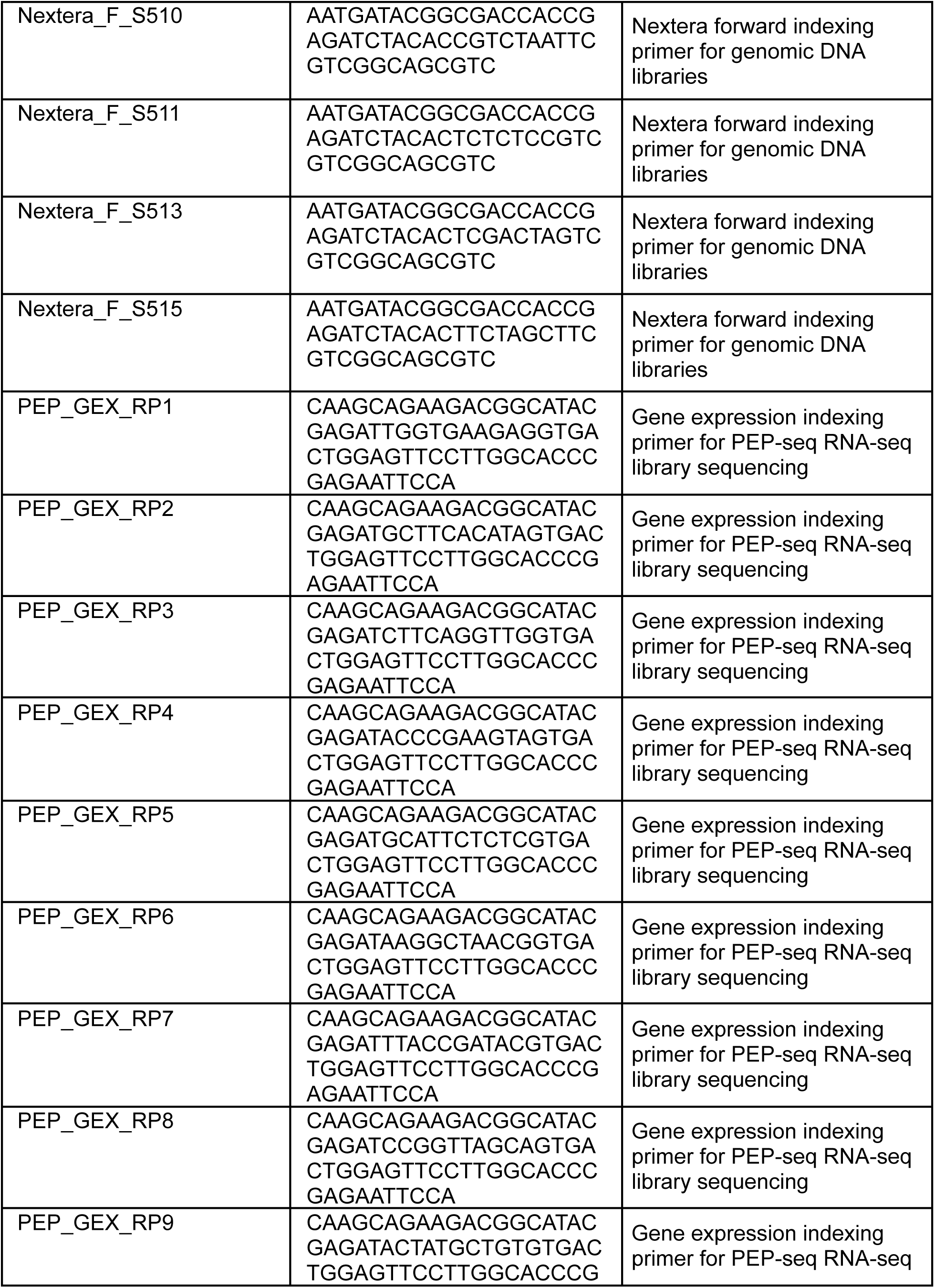
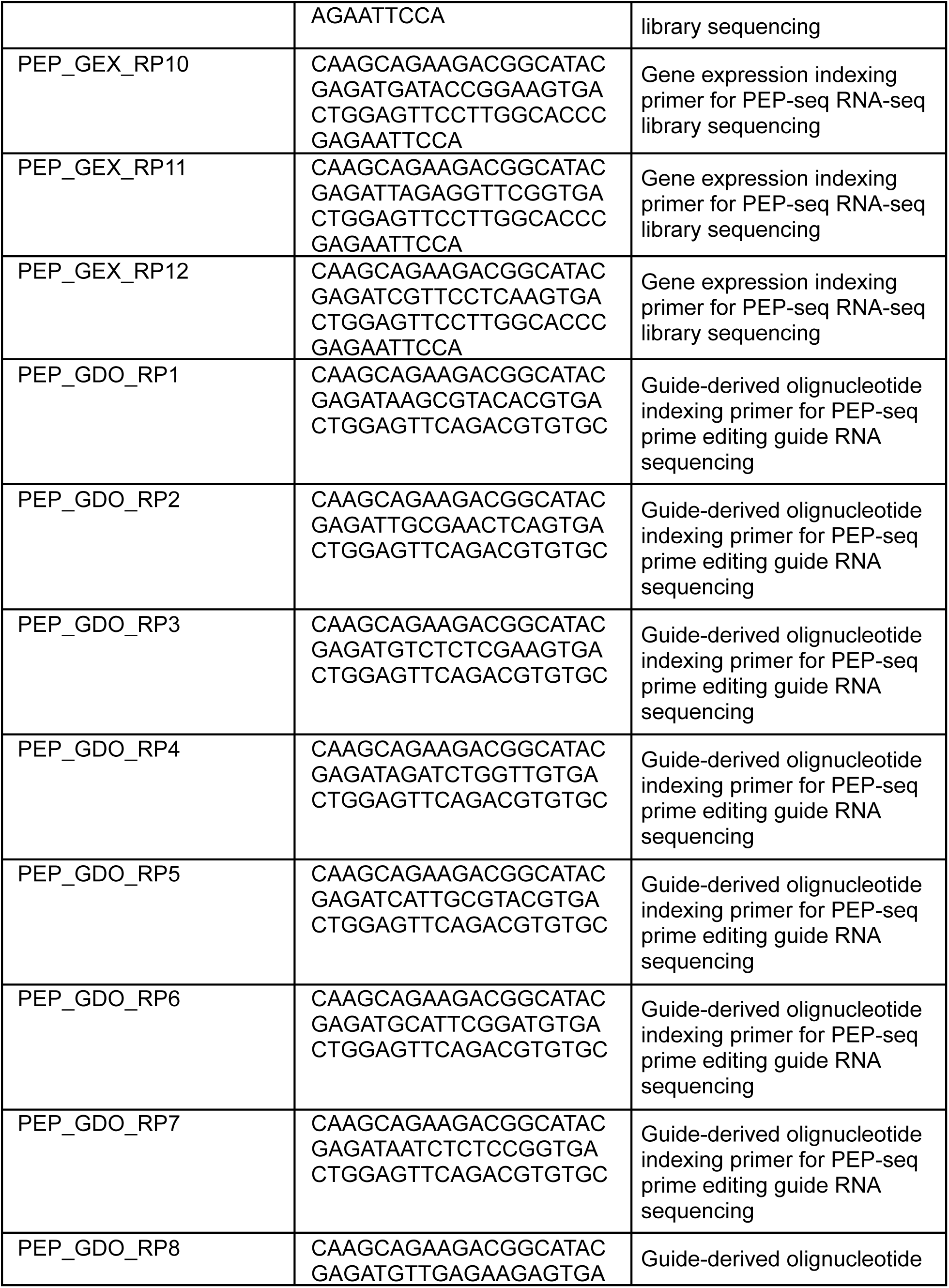
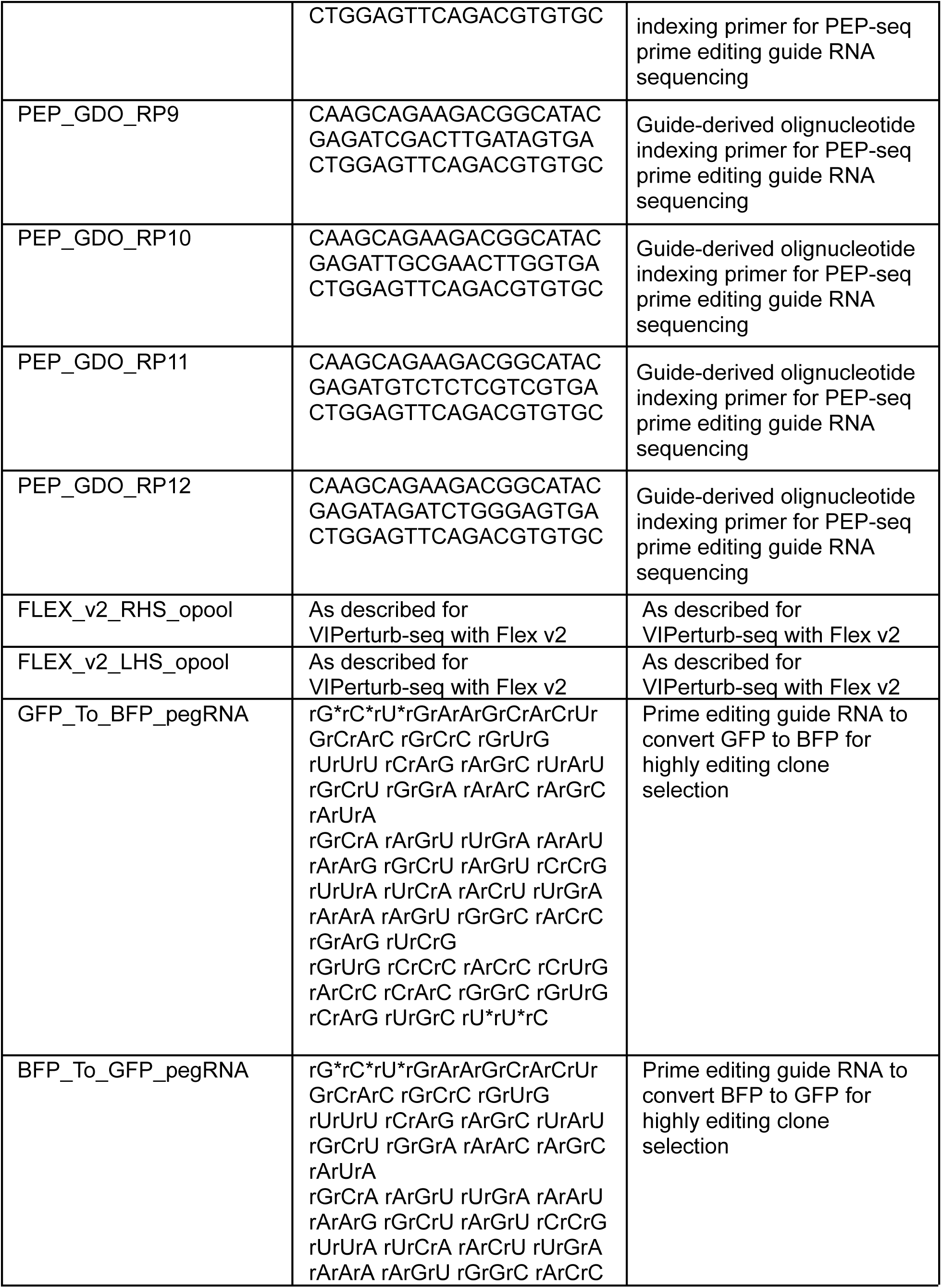
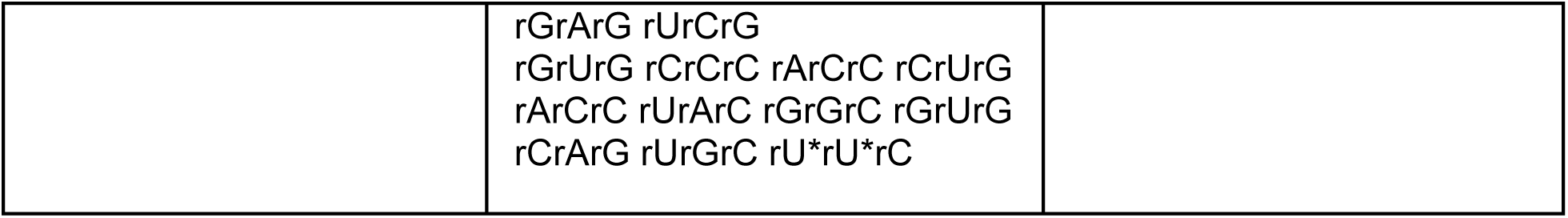
Oligonucleotide sequences, all were orders from Integrated DNA Technologies.

**Supplementary Table 2.**

| Reagent | Vendor | Cat number |
| --- | --- | --- |
| DMEM+GlutaMAX | ThermoFisher Scientific | 31966-047 |
| Fetal Bovine Serum | Sigma-Aldrich | F7524-500ML |
| Antibiotic-antimycotic | Corning | 30-004-CI |
| TrypLE™ Express Enzyme (1X), no phenol red | Gibco | 12604013 |
| 0.05% Trypsin=EDTA (1X) | Gibco | 25300-054 |
| Acridine Orange/Propidium Iodide (AO/PI) staining | Logos Biosystems | F23001 |
| Polybrene | VectorBuilder | PL0001 |
| Blasticidin S HCl | ThermoFisher Scientific | R21001 |
| SF Cell Line<br>4D-Nucleofector® LV Kit L | Lonza | V4LC-2002 |
| Puromycin Dihydrochloride | Gibco | A1113803 |
| GEM-X Flex Sample Preparation v2 Kit | 10X Genomics | PN-1000781 |
| GEM-X Flex GEM & Library Kit | 10X Genomics | PN-1000782 |
| GEM-X Flex Hybridization & Wash Kit | 10X Genomics | PN-1000789 |
| GEM-X Flex Human Transcriptome Probe Kit | 10X Genomics | PN-1000785 |
| GEM-X Flex Gene Expression Chip Kit | 10X Genomics | PN-1000791 |
| Dual Index Kit TS Set A, 96 | 10X Genomics | PN-1000251 |
| rxns |  |  |
| Allprep DNA/RNA kit | Qiagen | 80204 |
| Qubit 1X dsDNA HS Assay Kit | ThermoFisher Scientific | Q33230 |
| Qubit RNA High Sensitivity (HS) Assay Kit | ThermoFisher Scientific | Q32852 |
| NEBNext® Ultra™ II Q5® Master Mix | New England Biolabs | M0544S |
| SPRIselect DNA Size Selection Reagent, 60 mL | Beckman Coulter | B23318 |
| Diagenode Bulk ATAC-seq kit | Diagenode | C01080001 |

